# Evolutionary transitions from female to hermaphrodite reproduction remodel olfactory and mating-receptive behaviors

**DOI:** 10.1101/2023.10.16.562407

**Authors:** Margaret S. Ebert, Cornelia I. Bargmann

**Affiliations:** The Rockefeller University, 1230 York Avenue, New York, NY, USA

## Abstract

Male/hermaphrodite species have arisen multiple times from a male/female ancestral state in nematodes, providing a model to study behavioral adaptations to different reproductive strategies. Here we examined the mating behaviors of male/female (gonochoristic) *Caenorhabditis* species in comparison to male/hermaphrodite (androdiecious) close relatives. We find that females from two species in the *Elegans* group chemotax to volatile odor from males, a behavior described in only a few animal species. The females also display known mating-receptive behaviors such as sedation when male reproductive structures contact the vulva. Focusing on the male/female species *C. nigoni,* we show that female chemotaxis to males is limited to adult females approaching adult or near-adult males, and relies upon the AWA neuron-specific transcription factor ODR-7, as does male chemotaxis to female odor as previously shown in *C. elegans*. However, female receptivity during mating contact is *odr-7-*independent. All female behaviors are suppressed by mating, and all are absent in young hermaphrodites from the sister species *C. briggsae*. However, latent receptivity during mating contact can be uncovered in mutant or aged *C. briggsae* hermaphrodites that lack self-sperm. Young hermaphrodites from a second androdioecious species, *C. tropicalis*, are similarly unreceptive to males, but recover all female behaviors upon aging. These results reveal two mechanistically distinct components of the shift from female to hermaphrodite behavior: the loss of female-specific *odr-7-*dependent chemotaxis, and a sperm-dependent state of reduced receptivity to mating contact. The recovery of receptivity after sperm depletion has the potential to maximize hermaphrodite fitness across the lifespan.

**Highlights:**

- Female and hermaphrodite mating behaviors differ in closely related nematode species
- Females are attracted to volatile male odors, but hermaphrodites are not
- The same olfactory neuron pair drives female attraction to males and vice versa
- Latent female mating behaviors are revealed in hermaphrodites that lack self-sperm

## Introduction

In sexually reproducing species, innate mating behaviors evolve in concert with ecological and physiological adaptations. An extreme form of sexual adaptation has occurred within the *Caenorhabditis* clade of nematodes, where at least three lineages have independently evolved self-fertilizing hermaphrodites from the female sex of ancestral male/female species, giving rise to *C. elegans*, *C. briggsae*, and *C. tropicalis*.^1,2^ In all of these species, hermaphrodites produce a limited supply of sperm in the last larval stage, then produce oocytes throughout adulthood. Hermaphrodites can either self-fertilize to produce hermaphrodite progeny, or mate with rare males to produce male and hermaphrodite cross-progeny. The repeated appearance of androdioecy provides an opportunity to examine evolutionary transitions between ancestral female and derived hermaphrodite mating behaviors.

The mating behaviors of male nematodes have been characterized in *C. elegans* and a few other *Caenorhabditis* species,^3,4,5,6,7,8,9^ whereas the more limited analysis of female mating behavior has emerged primarily from studies of the gonochoristic species *Caenorhabditis remanei*. Virgin females of *C. remanei* approach conspecific male-female mating pairs^10^ via an attractant that has airborne transmission across a ∼1 mm distance^11^. Chemotaxis in a microfluidic arena shows that *C. remanei* females are attracted to non-mating males as well, likely via nonvolatile pheromones.^12^ When a *C. remanei* male presses his tail against the female’s vulva, a soporific factor requiring the male somatic gonad induces locomotor quiescence and rhythmic vulval muscle contractions in the female as male sperm is transferred.^10^ Several behaviors in *C. remanei* females are decreased after recent mating: mate search (leaving a food lawn in the absence of males),^13^ attraction to male odor,^11,12^ and male-triggered sedation.^10^

In contrast to *C. remanei* females, young adult *C. elegans* hermaphrodites do not leave food to seek mates^13^ and are not attracted to males.^12,14^ They resist male mating attempts, sprinting forward when males contact the vulva^10,15^ and keeping the vulva tightly closed against prodding by male partners.^10^ *C. elegans* hermaphrodites that are sperm-depleted by aging or by genetic mutation are more easily fertilized by males, ^10,15^ indicating that hermaphrodites may retain some female-like mating responses. Moreover, both aged hermaphrodites and *fog-2* mutants lacking self-sperm are more attractive to males than young adult hermaphrodites.^16,17,18^ However, interpretation of these results is limited by the few nematode species in which hermaphrodite or female behavior has been examined.

Within the *Caenorhabditis* genus, some closely related sister-species pairs differ in their reproductive modes, as for example in hermaphrodite *C. briggsae* versus female *C. nigoni*, and hermaphrodite *C. tropicalis* versus female *C. wallacei*.^2^ Here, we characterize female and hermaphrodite mating behaviors in these species and probe how the behaviors are modulated by the identity of the male partner, by developmental stage, and by mating experience. Taking advantage of advances in CRISPR technology^19,20^ and the genome sequence of *C. nigoni*,^21,22^ we provide insight into genetic mechanisms for female attraction to males. Using interspecific crosses, mutant analysis, and aging, we expose latent female behaviors in *C. briggsae* and *C. tropicalis* hermaphrodites and shed light on their regulation.

## Results

### Female nematodes display three mating behaviors: chemotaxis to males, sedation, and vulva opening

Three androdiecious species within the *Caenorhabditis* genus – *C. elegans, C. briggsae,* and *C. tropicalis* -- fall within the *Elegans* group, which also contains numerous gonochoristic species.^21,23,24^ To compare female and hermaphrodite mating behaviors, we examined the closely related sister-species pairs *C. nigoni* and *C. briggsae*, and *C. wallacei* and *C. tropicalis* (Figure 1A). *C. inopinata*, the recently discovered sister species to *C. elegans*, did not grow well under standard laboratory conditions, so it was not studied here.^25^ We first observed female mating behaviors in video recordings of wild-type virgin *C. wallacei* and *C. nigoni* mating pairs. As in *C. elegans*, the male scans the female’s body with the ventral side of his tail, making sharp turns at her head or tail, until he locates the vulva. During this initial scanning phase, the female makes small forward and reverse movements (Figure 1B, Video S1). At the moment that the male’s ventral tail touches her vulva, her locomotion ceases, her body posture relaxes, and her pharyngeal pumping slows, as described in *C. remanei*^10^ (Videos S1 and S2). She remains sedated for about one minute or more while the male attaches to her, protracts his spicules into the vulva, transfers sperm into the uterus, and deposits a mating plug onto the vulva. During this process, the female’s vulval muscles coordinately contract, widening the vulval slit (Figure 1C, Video S2).

**Figure 1.**
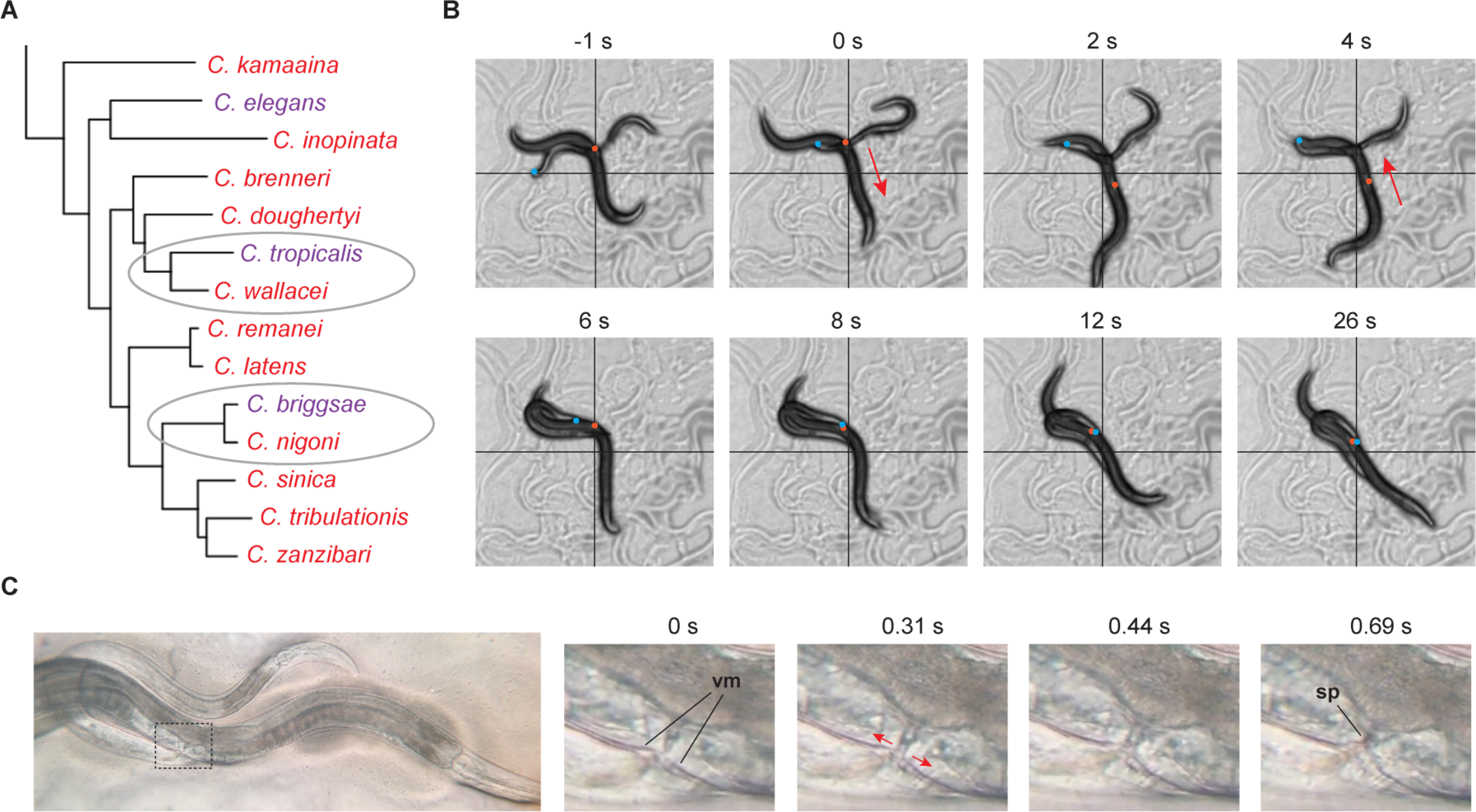
*Caenorhabditis* females perform receptive behaviors during mating. (A) Phylogeny of the *Elegans* group of *Caenorhabditis* species. Red, gonochoristic species with male and female sexes. Purple, androdiecious species with self-fertilizing hermaphrodites and cross-fertilizing males. Adapted from ^24^. (B) Female sedation during mating. *C. wallacei* male-female pair. t=0 denotes first male tail contact with the female. Red dot, female vulval slit; blue dot, male ventral tail. Arrows in the first four frames show direction of female movement. See also Videos S1-S3. (C) Female vulva opening during mating. *C. nigoni* male-female pair. t=0 denotes first male spicule prodding of the vulva. Right panels, inset from dashed rectangle. vm, vulval muscles; sp, spicules. Red arrows show direction of vulval muscle contraction. See also Video S2. Females in panels B and C are 1 mm long.

In contrast with the females from their sister species, young adult hermaphrodite *C. briggsae* or *C. tropicalis* moved vigorously to escape male contact at the vulva when paired either with males of their own species or with males of closely related gonochoristic species (Table 1, Video S3). Similarly, vulva opening was not observed in young adult *C. briggsae* or *C. tropicalis* hermaphrodites, either with males of the same species or with gonochoristic male partners (Table 1, Video S3). In subsequent experiments, *C. nigoni* and *C. wallacei* males were used because *C. briggsae* ^10^ and *C. tropicalis* males were unreliable mating partners (like *C. elegans* males^10^). For example, only 1/10 *C. tropicalis* males versus 6/10 *C. wallacei* males prodded the vulva of a *C. tropicalis* hermaphrodite in five-minute assays. By contrast, males from male-female species make vigorous mating attempts to females and hermaphrodites.^26^

**Table 1.**
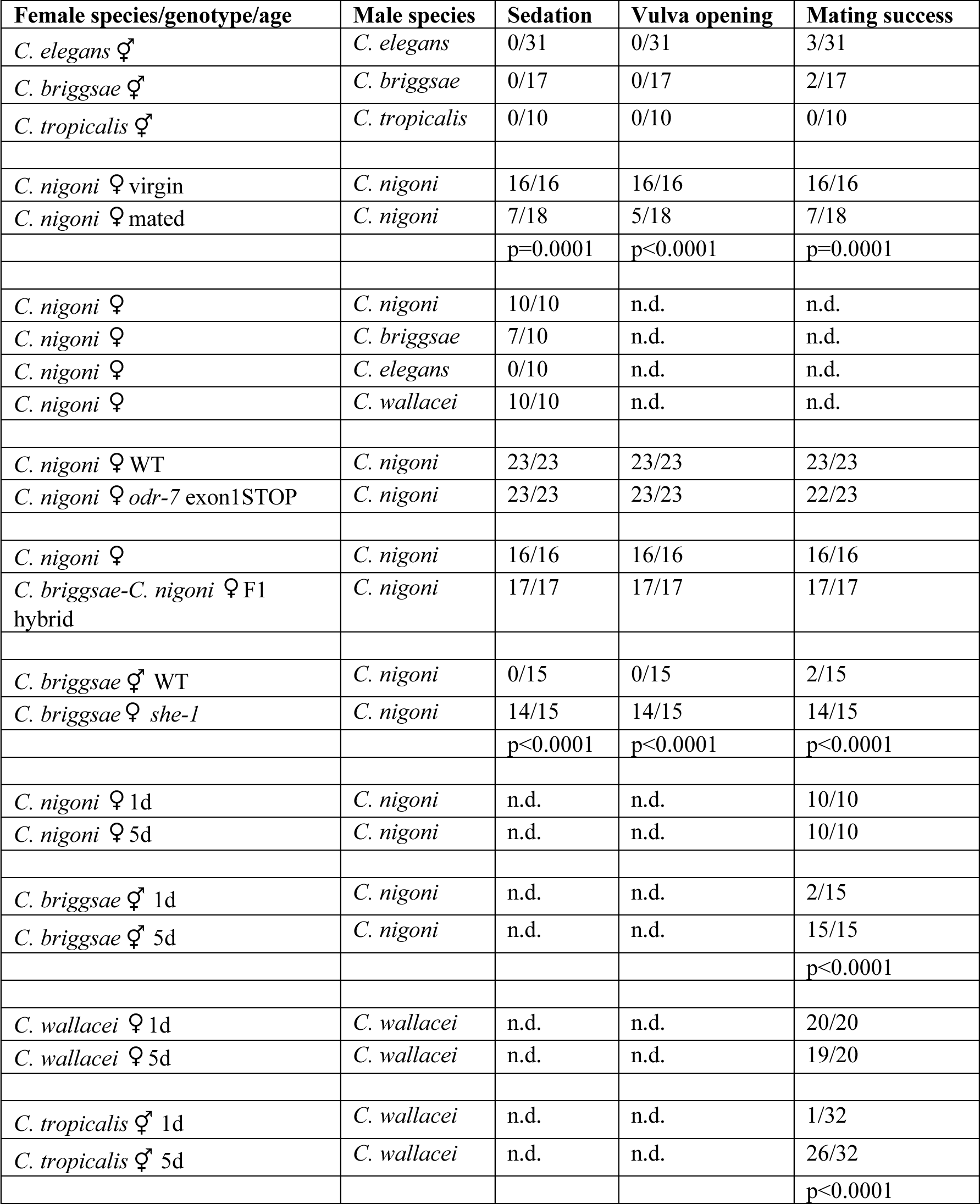
Female/hermaphrodite receptive behaviors. Behavioral outcomes scored for five-minute assays. Pairwise comparisons: Fisher’s exact test with two-tailed p-value.

Direct observation of the mating assays suggested that *C. nigoni* or *C. wallacei* females were attracted to male partners. When 20 *C. nigoni* or *C. wallacei* virgin females were placed on a plate with a single conspecific male, the females moved rapidly toward the male and any female mating partner (Figure 2A, Figure S1A, Video S4). By contrast, *C. briggsae* and *C. tropicalis* young adult hermaphrodites did not approach conspecific males or gonochoristic sister species males (n ≥ 10 assays). We simplified the chemotaxis assay by using individual roller mutant *C. nigoni* males generated by CRISPR-Cas9 mutagenesis; a roller animal moves in a tight circle, effectively acting as a point source of odor. Roller *C. nigoni* males attracted as many *C. nigoni* females as wild-type males (Figures 2C, S1B, and S2F), but roller females did not attract females (Figure 2E), establishing the sex-specific nature of the attractant. In a similar chemotaxis assay with 20 males and one female, males moved quickly to the single roller female (Figures 2B and 2D, Video S5) but not to a single roller male (Figure 2F).

**Figure 2.**
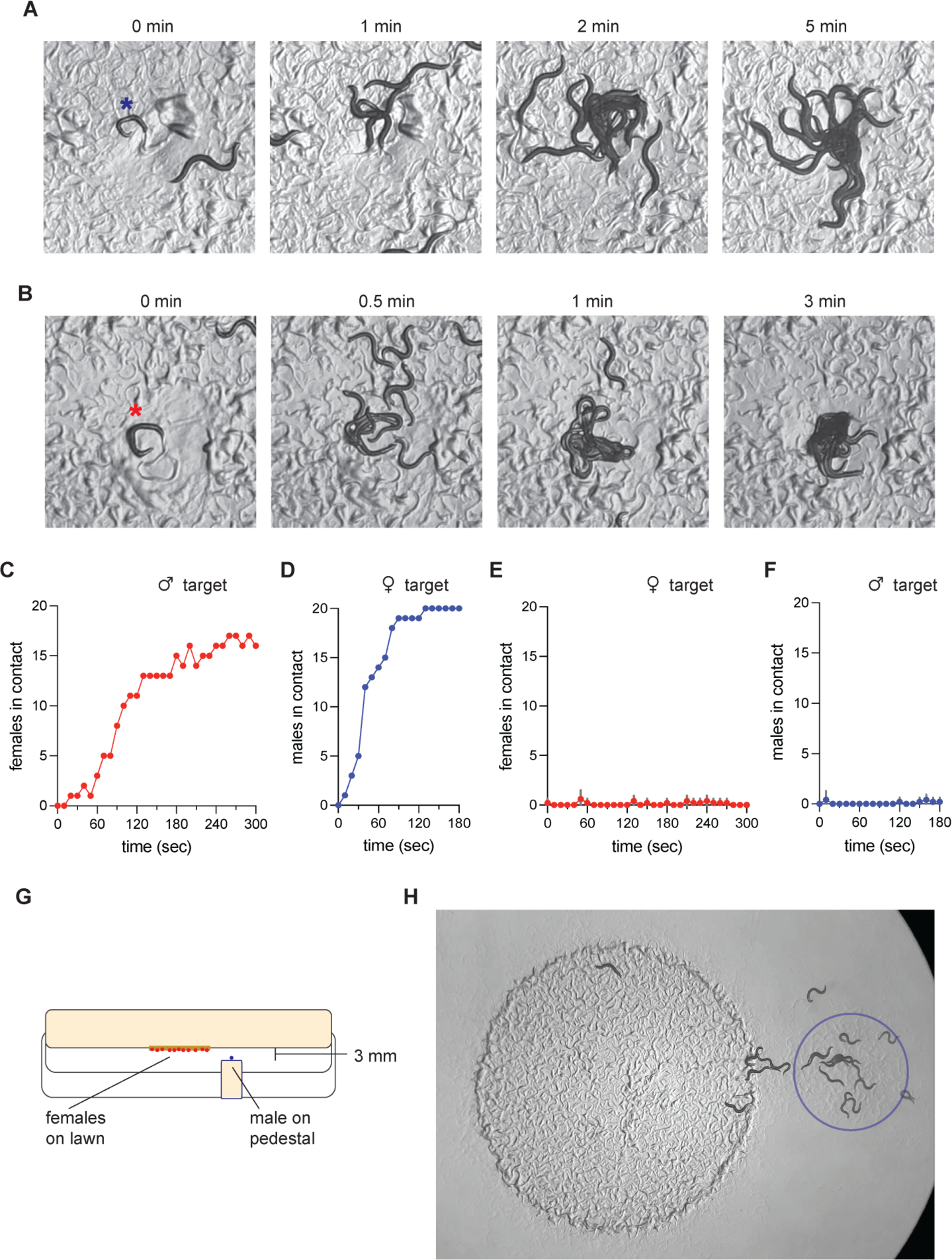
*C. nigoni* females chemotax to a volatile male odor. (A-B) Representative images from *C. nigoni* mate attraction assays. One roller target animal was placed on a lawn with 20 test animals at t=0. Time of image is noted. (A) Male target (blue asterisk) with 20 virgin females. See also Video S4. (B) Female target (red asterisk) with 20 virgin males. See also Video S5. (C-F) Mate attraction assays, showing number of test animals in contact with the target animal or each other. (C) Male target with 20 virgin females, representative assay. See also Video S4. (D) Female target with 20 virgin males, representative assay. See also Video S5. (E) Female target with 20 virgin females (n = 5 assays each). (F) Male target with 20 virgin males (n = 5 assays each). Red dots (E) or blue dots (F) show mean at each timepoint; gray error bars, +/-SD. (G) Diagram of the pedestal attraction assay. A roller *C. nigoni* target male was placed on an agar pedestal 3 mm below the agar surface of a plate with 20 *C. nigoni* virgin females. (H) Still image of a representative pedestal assay after 5 minutes. Blue circle, position of the pedestal. Females in panels A and H are 1 mm long. Males in panel B are 0.8 mm long. See also Figure S1 and Supplementary Data Table 2H.

To distinguish whether females are attracted to males directly or indirectly via attraction to a mating pair, we examined assays with one virgin *C. nigoni* female paired with one roller *C. nigoni* male (Figure S1C). The dynamics of a single female’s approach to the male were indistinguishable from the dynamics in assays with 20 females (Figures S1B-C), indicating that individual females chemotax efficiently to a male odor with no requirement for the male to be mating with another female.

The rapid movement of females toward the male suggested that the male attractive odor could be volatile. To examine this possibility, we assayed the behavior of *C. nigoni* females in response to a roller male placed on a small agar pedestal ∼3 mm below the lawn (Figure 2G). Females moved toward the region above the male pedestal with similar dynamics as when placed on the same agar surface (Figure 2H, Supplementary Data Table 2H). Indeed, females would exit a bacterial food lawn to approach the male pedestal. Thus, females can sense and chemotax to a volatile male odor that diffuses through air.

### Developmental stage, mating status, and species of mating partner affect female mating behavior

To determine when developing animals first emit and respond to attractive odors, we paired *C. nigoni* males and females at different stages. Individual L4 males elicited a range of chemotaxis responses from adult females (Figure 3A), suggesting that *C. nigoni* males begin to emit attractive odor in the last larval stage, before full sexual maturity. By contrast, L4 larval stage *C. nigoni* females were not attracted to an adult roller *C. nigoni* male at all (Figure 3B), although they moved actively around the lawn. Thus, females do not respond to male odor until they are sexually mature.

**Figure 3.**
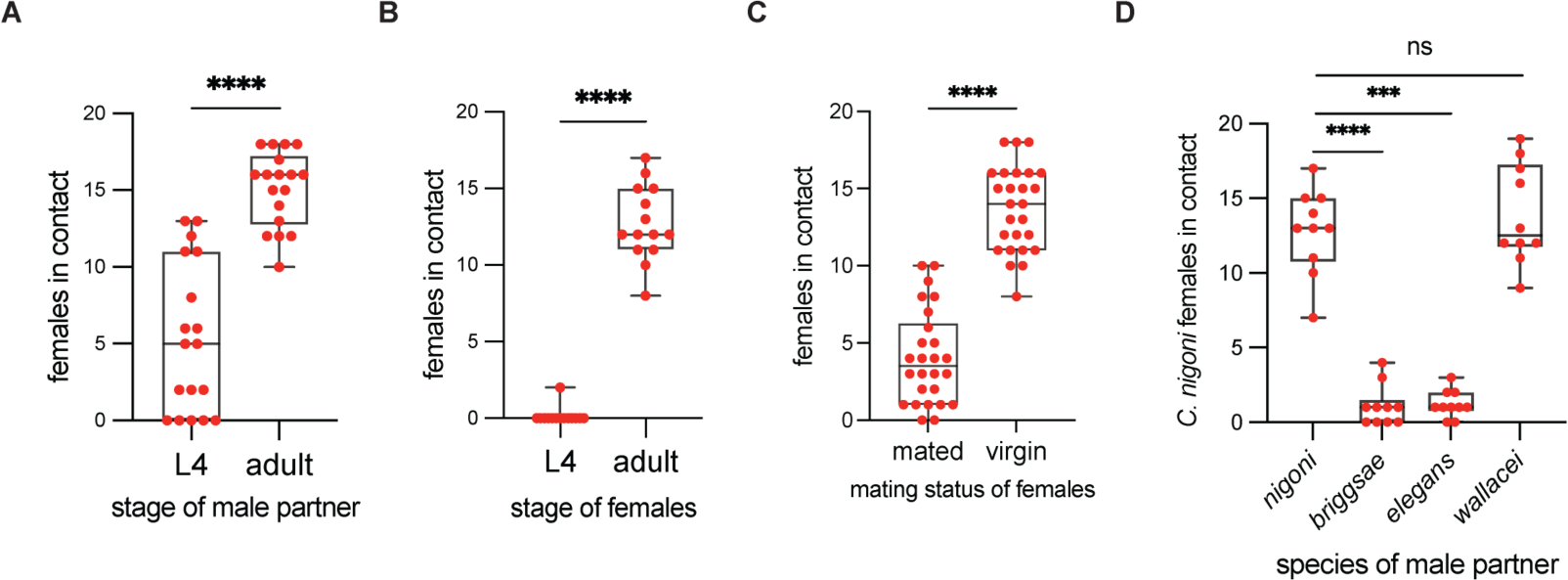
Attraction to males is modulated by developmental stage and mating history. (A-D) Chemotaxis assays with 20 *C. nigoni* females and one target male, scored at 5 minutes. (A) Adult female chemotaxis to L4 larval stage or 1-day adult roller males (n = 18 assays each). (B) Chemotaxis of L4 larval stage or 1-day adult females to adult roller male (n = 14 assays each). (C) Chemotaxis of recently mated or virgin females to adult roller male (n = 26 assays each). See also Figure S3C. (D) *C. nigoni* female chemotaxis to wild-type males of different species (n = 10 assays each). Center bar and box denote median and quartiles across assays; flanking bars denote full range. Pairwise comparisons: Mann-Whitney test. Three-way comparisons: Kruskal-Wallace test with Dunn’s multiple comparisons test. ns P>0.05, *** P<0.001, **** P<0.0001. See also Table 1.

Female behavior in many animals is altered after mating. We prepared batches of recently mated females by breeding *C. nigoni* virgin females to wild-type *C. nigoni* males, and then assayed their behavior within four hours of mating. Recently mated females showed significantly less chemotaxis to a roller *C. nigoni* male (Figure 3C), and less sedation and less vulva opening than virgin females of the same age (Table 1). Virgin females that were pre-exposed to males but not mated were still attracted to males (Figure S3C).

Female discrimination between potential mating partners underlies reproductive isolation in many animals.^27^ We examined the response of *C. nigoni* females to individual males from four different species: conspecific *C. nigoni*, the androdiecious sister species *C. briggsae*, the more distantly related androdiecious *C. elegans*, or the more distantly related gonochoristic *C. wallacei*. *C. briggsae* and *C. elegans* males did not attract *C. nigoni* females (Figure 3D), but *C. wallacei* males were as attractive to *C. nigoni* females as conspecific males (Figure 3D). Moreover, *C. nigoni* females were sedated by *C. nigoni* and *C. wallacei* males with equal success, and additionally sedated by seven of ten *C. briggsae* males; *C. elegans* males failed to sedate any *C. nigoni* female partner (Table 1). Together, these results demonstrate unexpected attraction and receptivity of *C. nigoni* females to males of several species.

### Sex-specific attraction behaviors in *C. nigoni* require the shared transcription factor ODR-7

In *C. elegans*, rapid male chemotaxis to volatile female odor requires the nuclear hormone receptor transcription factor ODR-7.^18^ To test whether ODR-7 functions in mate attraction in *C. nigoni*, we first confirmed that the *odr-7* gene is present in *C. nigoni* and identified two *odr-7* mRNA isoforms by reverse transcription-PCR (Figure 4A, Supplemental Information). Both isoforms are expected to encode functional ODR-7 protein including a predicted DNA-binding domain at the C-terminus.

**Figure 4.**
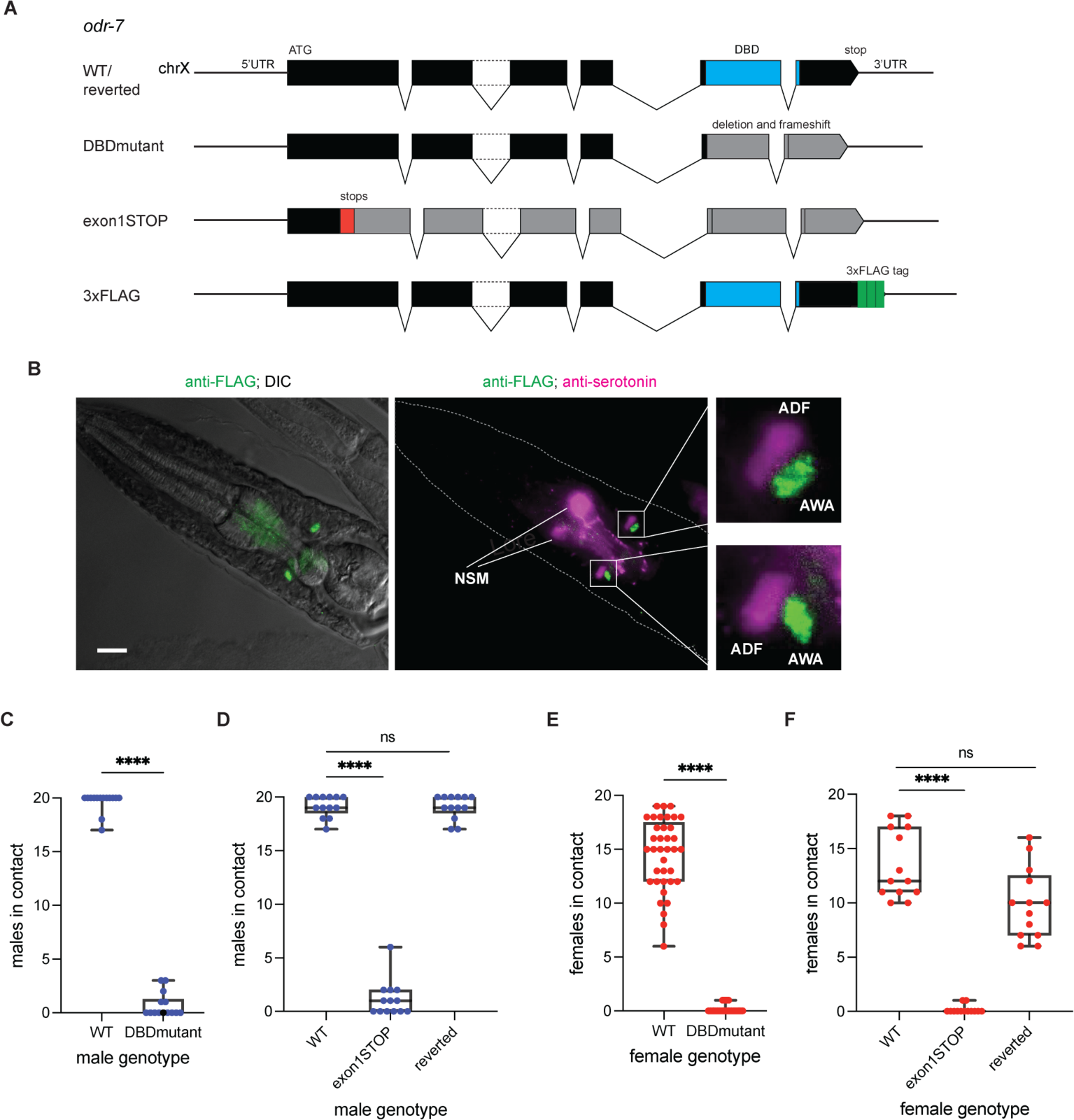
*odr-7* is required for male-to-female and female-to-male *C. nigoni* chemotaxis. (A) Structure of wild-type and mutant *odr-7* genes. Two mRNA splice forms remove intron 2 or retain it as coding sequence (dotted black lines). Black rectangles, exons. Blue, predicted DNA binding domain (DBD). Gray, frameshifted sequence downstream of CRISPR mutation. Red, 43-bp STOP-IN CRISPR cassette insertion. Green, 66-bp 3xFLAG tag. (B) ODR-7::FLAG expression in *C. nigoni*. (Left) Adult *odr-7*::3xFLAG male immunostained with anti-FLAG antibody (green) superimposed on DIC image. (Center) The same *odr-7*::3xFLAG male showing immunostaining for FLAG (green) and serotonin, which is expected to label ADF sensory neurons and NSM pharyngeal neurons (magenta). (Right) Magnified images in the region of AWA and ADF. Scale bar, 10 *μ*m. Anterior is at left. Note background anti-FLAG staining in the pharynx; see also Figures S2C-E. (C-F) Chemotaxis assays with 20 *C. nigoni* test animals and one roller target animal. (C) Chemotaxis of wild type or *odr-7* DBD mutant males to female target, scored at three minutes (n = 14 assays each). (D) Chemotaxis of wild type, *odr-7* exon 1 STOP-IN mutant, or reverted exon 1 *odr-7* males to female target; scored at three minutes (n = 13 assays each). (E) Chemotaxis of wild-type or *odr-7* DBD mutant females to male target, scored at five minutes (n = 37-42 assays each). See also Figure S3C. (F) Chemotaxis of wild-type, *odr-7* exon 1 STOP-IN mutant, or reverted exon 1 *odr-7* females to male target; scored at five minutes (n = 13 assays each). See also Figures S2F-G, Table 1. Pairwise comparisons: Mann-Whitney test. Three-way comparisons: Kruskal-Wallace test with Dunn’s multiple comparisons test. **** P<0.0001.

In *C. elegans*, *odr-7* expression is specific to the bilateral AWA olfactory neurons in the amphid sensory organs.^28^ Neuronal positions and identities are often conserved between different *Caenorhabditis* species,^29^ but some variation in sensory cell types has been observed in more distant *Strongyloides* and *Pristionchus* nematodes.^29,30,31^ We confirmed that four pairs of amphid sensory neurons, identified by staining with the vital dye DiO, were present in the same locations in *C. nigoni* as in *C. elegans* (Figure S2A). To see where *odr-7* was expressed in *C. nigoni,* we epitope-tagged the endogenous *odr-7* by inserting a 3xFLAG tag^32^ at the extreme C-terminus using CRISPR mutagenesis (Figure 4A). Both males and females of the epitope-tagged *C. nigoni* strain had wild-type chemotaxis to the opposite sex (Figure S2B). Immunostaining showed that ODR-7 protein was expressed robustly and exclusively in the nuclei of a single neuron pair in both males and females (Figure 4B, S2C-E). The staining was in the expected location of the AWA neuron pair based on its position and its adjacency to a sensory neuron pair that contains serotonin (Figure 4B, S2C-D); serotonin is made by the ADF neuron that resides immediately anterior to AWA in *C. elegans*.^33^ These results suggest that *odr-7* is expressed exclusively in AWA neurons of adult *C. nigoni* females and males.

We used CRISPR-Cas9 mutagenesis to generate a predicted null mutant of *odr-7* in *C. nigoni* by deleting 43 bp at the beginning of the highly conserved DNA-binding domain (DBD) (“DBDmutant,” Figure 4A).^34^ Males from the *odr-7* DBD mutant did not chemotax to a roller female (Figure 4C), recapitulating the chemotaxis defect in *C. elegans odr-7* males.^18^ A second *C. nigoni odr-7* mutant was made by CRISPR insertion of a cassette that introduces stop codons and a frameshift into the first exon (“exon1STOP,” Figure 4A).^20^ The *odr-7* exon1STOP mutant was also defective in male chemotaxis to females (Figure 4D). When the STOP-IN cassette was deleted in a second CRISPR mutagenesis (“reverted,” Figure 4A), males of the resulting strain were fully rescued for chemotaxis to females (Figure 4D). A different STOP-IN mutation did not cause a mutant phenotype, likely due to translational re-initiation (Figure S3A-B, Supplemental Information).

Strikingly, *odr-7* was also required for female *C. nigoni* chemotaxis to males. In contrast with wild-type *C. nigoni* females, *odr-7* DBD mutant or exon1STOP mutant females did not approach roller males in a chemotaxis assay (Figure 4E-F). The reverted *odr-7* exon 1 strain resembled the wild-type strain, confirming the causal role of *odr-7* (Figure 4F). *odr-7* mutant females were defective in chemotaxis both to roller and to wild-type male partners (Figure S2F) and remained defective when induced into an aroused locomotor state by tapping the tail (Figure S2G). However, *odr-7* mutant females had normal male-triggered sedation and vulva opening during mating (Table 1), separating female chemotaxis from the other components of female mating behavior.

### Latent female behavior in *C. briggsae* and *C. tropicalis* hermaphrodites

What are the barriers to female behavior in androdiecious species? The closely related species *C. nigoni* and *C. briggsae* can interbreed to form hybrid F1 offspring that develop as females with no self-sperm.^35^ Virgin *C. nigoni*/*C. briggsae* F1 females were largely attracted to a *C. nigoni* male (Figure 5A), resembling *C. nigoni* females rather than *C. briggsae* hermaphrodites. F1 hybrid females also responded robustly to *C. nigoni* male partners with the receptive behaviors of sedation and vulva opening (Table 1). Thus female behaviors appeared dominant to those of hermaphrodites in the F1 hybrids. Using these F1 hybrids, we asked whether changes in *odr-7* account for the absence of *C. briggsae* hermaphrodite chemotaxis to males. F1 hybrid females between *C. nigoni odr-7* exon1STOP mutants and wild-type *C. briggsae*, whose only intact copy of *odr-7* was inherited from *C. briggsae*, were proficient in chemotaxis to *C. nigoni* males (Figure 5A), suggesting that the lack of chemotaxis in *C. briggsae* does not result from a defective *odr-7* gene. Instead, it could result either from other sensory changes or from the presence of sperm in hermaphrodites.

**Figure 5.**
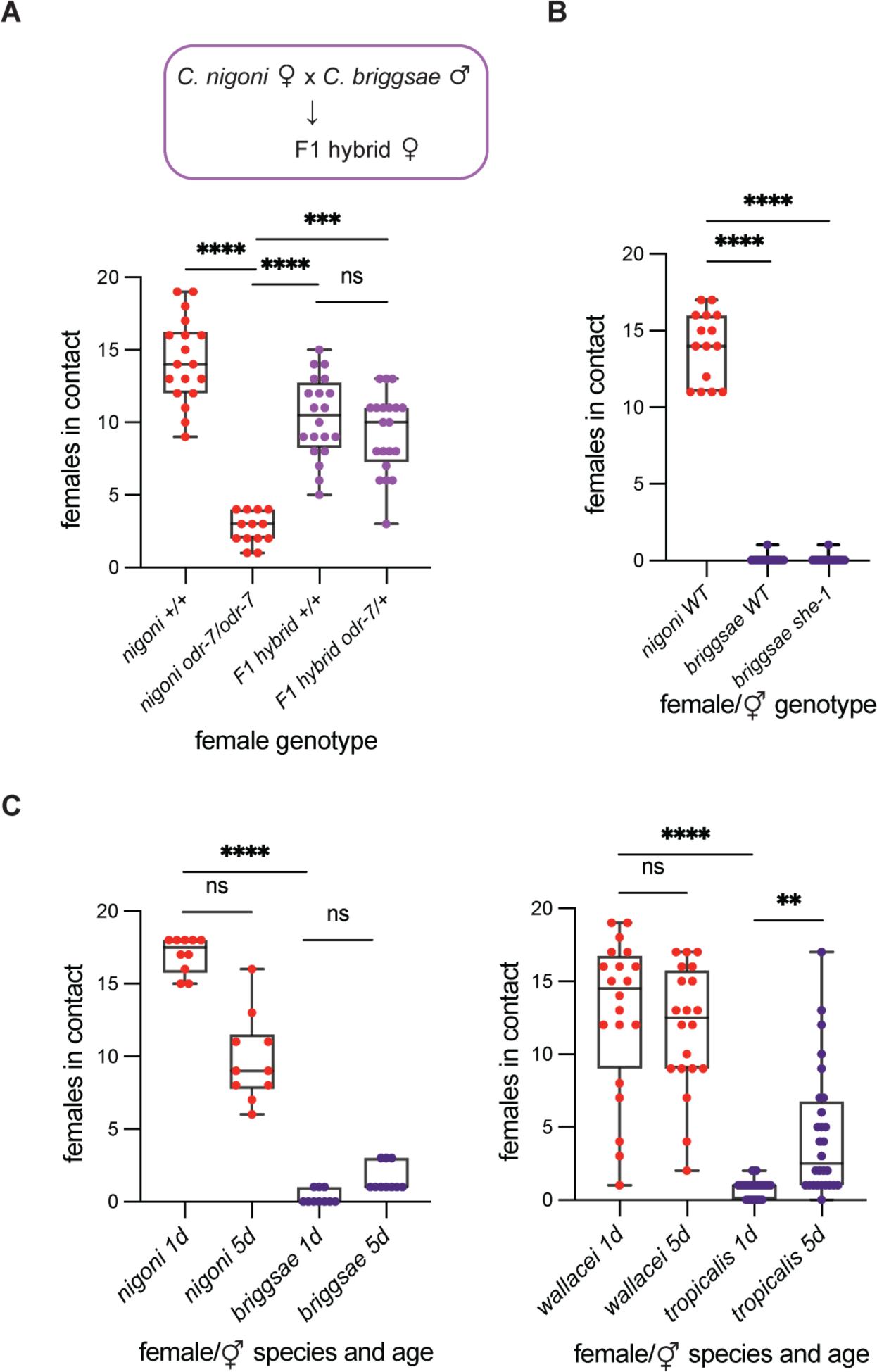
Latent female behaviors in *Caenorhabditis* hermaphrodites. (A) Chemotaxis of *C. nigoni* females or F1 hybrid *C. nigoni/C. briggsae* females to wild-type *C. nigoni* males, scored for maximum females in contact during 5-minute assay (n = 20 assays each). Hybrids were F1 progeny of *C. briggsae* males and *C. nigoni* wild-type or *odr-7* exon 1 STOP-IN mutant females. (B) Chemotaxis of *C. nigoni* females, *C. briggsae* hermaphrodites, or *C. briggsae she-1* mutant pseudofemales to roller *C. nigoni* male target; scored at five minutes (n = 15 assays each). See also Video S6. (C) Chemotaxis of 1-day or 5-day old *C. nigoni* females or *C. briggsae* hermaphrodites to a wild-type *C. nigoni* male (left, n = 10 assays each) or *C. wallacei* females or *C. tropicalis* hermaphrodites to a wild-type *C. wallacei* male (right, n = 20-30 assays each). At 5 days, hermaphrodites are depleted of self-sperm. All assays are groups of 20 females with one target male. (B-C) are scored at five minutes. Three- and four-way comparisons: Kruskal-Wallace test with Dunn’s multiple comparisons test. ns P>0.05, ** P<0.01, *** P<0.001, **** P<0.0001. See also Table 1.

To probe the role of sperm in regulating hermaphrodite mating in *C. briggsae*, we used a temperature-sensitive mutant strain (*she-1*) wherein XX animals grown at a non-permissive temperature develop eggs but no self-sperm (“pseudofemales”).^36^ These *she-1 C. briggsae* pseudofemales did not chemotax to males, resembling wild-type *C. briggsae* hermaphrodites (Figure 5B). However, *C. briggsae she-1* pseudofemales showed robust sedation and vulva opening upon male contact, like *C. nigoni* females (Table 1, Video S6). This result suggests that self-sperm or fertilized eggs in young adult *C. briggsae* hermaphrodites suppress female receptive behaviors triggered by vulval contact.

In hermaphrodites, sperm are produced in limited numbers prior to prolonged oocyte production. Like *she-1 C. briggsae* mutants, aged, sperm-depleted *C. briggsae* hermaphrodites were receptive to immediate spicule insertion and sperm transfer by males (Table 1) but did not chemotax to a *C. nigoni* male (Figure 5C). Similarly, aged *C. tropicalis* hermaphrodites permitted rapid spicule insertion and sperm transfer by the gonochoristic sister species *C. wallacei* (Table 1). Aged *C. tropicalis* hermaphrodites also recovered partial chemotaxis to males (Figure 5C). Thus, aged hermaphrodites can exhibit latent mating receptivity as well as female-like attraction to males, depending on the species.

## Discussion

By examining female and hermaphrodite behaviors in multiple *Caenorhabditis* species, including closely related species pairs with different reproductive strategies, we have identified reproducible female and female-like mating behaviors. We found that gonochoristic *C. nigoni* and *C. wallacei* females, but not androdioecious *C. briggsae* or *C. tropicalis* hermaphrodites, are strongly attracted to male volatile odors. Females and males of many species are robustly attracted to nonvolatile sex pheromones, and many female animals attract males at a distance by emitting volatile pheromones, but there are relatively few examples of female attraction to male volatile pheromones.^37,38,39,40,41,42^ Notably, these exceptions include two nematode species,^11,43^ as well as the two described here. *C. nigoni* males and females mutually send and respond to sex-specific volatile attractants, and chemotaxis in both sexes requires the AWA-specific transcription factor ODR-7, suggesting that female and male *C. nigoni* use the sex-shared AWA olfactory neurons to detect sex-specific attractants from potential mates. To our knowledge, this direct reprogramming of sex-shared olfactory neurons has not been seen in other species. Instead, for example, in silkworm moths the female pheromone bombykol is detected by different sensory neurons in males and females,^44^ and in *Drosophila*, both sexes express the odorant receptor that detects the volatile male sex pheromone cVA, but differences in higher brain circuits enable different responses for courtship behavior.^45^

As *C. briggsae odr-7* appears to complement a *C. nigoni odr-7* mutation in *C. briggsae/C. nigoni* hybrid females, the distinction between species may arise not from *odr-7* but from other genes. G protein coupled odorant receptors have undergone rapid evolution and diversification in animals, including *Caenorhabditis*.^46^ We speculate that differential expression of receptor genes in AWA of male and female *C. nigoni* allows differential mate attraction, and that the relevant genes are either absent or not expressed in AWA in *C. briggsae* hermaphrodites. This model aligns with the known age-, sex-, and experience-dependent sensory properties of the AWA olfactory neurons in *C. elegans.* Both male and hermaphrodite *C. elegans* require *odr-7* and AWA for chemotaxis to the food odor diacetyl;^28^ this behavior is stronger in hermaphrodites, which have higher AWA expression of the diacetyl receptor ODR-10.^13,47,48^ In males, the AWA neurons also detect attractive hermaphrodite odors^49^ due to sexually dimorphic expression of the G protein-coupled receptor SRD-1.^18^ AWA receptor gene expression is further modified by food deprivation.^48,50^ The model that AWA receptor expression contributes to the transition from female to hermaphrodite behaviors makes testable hypotheses for future studies of sex-specific receptor expression in *C. nigoni* and *C. briggsae*.

Female-like mating-receptive behaviors are mechanistically distinct from chemotaxis to males in *C. nigoni* and *C. briggsae: odr-7* mutant *C. nigoni* females, *she-1* mutant *C. briggsae* pseudofemales that lack sperm, and aged sperm-depleted *C. briggsae* hermaphrodites are all fully defective in chemotaxis to males, but fully proficient in contact-dependent receptive behaviors. Notably, receptive behaviors are regulated both in female *C. nigoni,* where they are suppressed by prior mating, and in hermaphrodite *C. briggsae*, where they are restored after aging. A similar suppression of mating-receptive behaviors is observed in *C. remanei* females after mating with wild-type but not germline-ablated males.^10^ Together, these results suggest that either sperm, seminal fluid, or fertilized eggs act as satiety signals for *Caenorhabditis* female receptive behaviors across both gonochoristic and androdioecious species. We speculate that mating behaviors in hermaphrodites are not merely degenerate female behaviors, but instead enhance their genetic fitness: young hermaphrodites minimize genetic dilution by avoiding mating while self-sperm are present, but can then increase total progeny number by allowing cross-fertilization after self-sperm are depleted. Although this strategy creates sexual conflict with males early in life, it enables other adaptive advantages of sexual reproduction.^26^

This study has several limitations. While we have increased the number of species in which female and hermaphrodite behavior has been examined, compared to prior work in *C. remanei* and *C. elegans,* additional species could have different strategies. We note also that only one wild-type strain was used to represent each species, and it is possible that female or hermaphrodite mating behaviors vary across wild strains.^51^ Mating behavior may also vary across environmental conditions, e.g. food source, starvation, or population density.^52^ We have found that *C. nigoni* does not express neuronal transgenes consistently even when using protocols optimized in other less-tractable nematode species such as *C. briggsae*^53^ or *Pristionchus pacificus*,^54^ and in the absence of efficient *C. nigoni* transgenesis, direct measurements or perturbations of AWA activity were not feasible. Finally, further experiments are required to define the sperm-related signal that suppresses female receptivity, and to ask whether the same signal suppresses chemotaxis to males in mated *C. nigoni* females and young adult *C. tropicalis* hermaphrodites. These are all avenues for further study.

## STAR Methods

### RESOURCE AVAILABILITY

#### Lead contact

Requests for strains, information or datasets should be directed to the lead contact, Cornelia I. Bargmann.

#### Materials availability

Strains generated in this study are available upon request to the lead contact.

#### Data and code availability

- All data reported in this paper are listed in Supplementary Tables. Sequence data have been deposited to GenBank.
- This paper does not report original code.

### EXPERIMENTAL MODEL DETAILS

#### Nematode strains

Animals were cultivated at 20-25°C on NGM agar plates using *E. coli* OP50 as the food source. The following strains were used for experiments (see Key Resources Table for details): *C. nigoni* wild-type JU1325, *rol-6(ky1036)* “roller,” *odr-7(ky1134)* “DBDmutant,” *odr-7(ky1100)* “exon1STOP,” *odr-7(ky1100ky1136)* “reverted,” *odr-7(ky1127)* “exon2STOP,” *odr-7(ky1135)* “3xFLAG;” *C. briggsae* wild-type AF16, *she-1(v35)*; *C. wallacei* wild-type JU1873; *C. tropicalis* wild-type JU1373; *C. elegans* wild-type PD1074. The *C. briggsae she-1* strain was always grown at 25°C to propagate as males and females. Gonochoristic strains were passaged with at least ∼100 individuals to avoid inbreeding bottlenecks.

#### Bacterial strains

*E. coli* OP50 was grown in LB liquid media at 20-25°C and seeded on standard NGM plates grown at 20-25°C.

### METHOD DETAILS

#### Behavioral assays

All animals used in mating behavior experiments were well-fed throughout life, and all assays took place on *E. coli* OP50 food lawns. Unless otherwise noted, males and females were 1-day virgin adults, separated by sex as L4 larvae one day before the assay. For L4 assays, L4 animals were picked the day of the assay. For 5-day adults, L4 animals were separated by sex five days before the assay. For mated females, 1-day virgin adults were bred to males, confirmed by the presence of a mating plug, and assayed within four hours of mating. Mating plates were seeded with 10 μl of *E. coli* OP50 diluted 1:3 from stationary liquid culture onto a 6 cm plate the day before the assay and grown at 20-25°C. Test animals (usually 20 females or males) were placed on a food lawn at least 30 minutes before placing a target animal on the lawn at t=0. For the pedestal assay, an agar cylinder was cut from a seeded NGM mating plate with a cork borer and placed on a plate lid; a roller target animal was placed on this pedestal, and then a plate with 20 females on a bacterial lawn was inverted over the pedestal/lid. All other mating and chemotaxis assays were performed on uncovered plates.

All chemotaxis assay data were collected on multiple days. For female chemotaxis, we found that groups of females could be tested serially with different males if we removed the previous male, replaced any mated female (recognizable by a mating plug) with a virgin female, and allowed the females at least 30 minutes to recover and redistribute over the lawn before adding a fresh male. The chemotaxis response was indistinguishable when assayed naively or after repeated trials (Figure S3C). Accordingly, we included data from up to ten repeats, but typically up to three repeats, in some assays (details in Supplementary Tables). Male groups were only tested once.

#### Chemotaxis quantification

##### Female chemotaxis

5-minute video recordings were manually scored at 10-second intervals for the number of females making any bodily contact with the male or with any female in the cluster with the male. For assays using wild-type males, t=0 is the moment the male begins scanning a female with the ventral side of his tail, typically within one minute of placing the male on the lawn. For assays using roller males, t=0 is the moment the male is placed on the lawn. *C. nigoni*/*C. briggsae* F1 hybrid female assays were quantified for the maximum cluster size within 5 minutes; all other assays were quantified at the five-minute endpoint.

##### Male chemotaxis

3-minute video recordings were manually scored at 10-second intervals for the number of males making any bodily contact with the female or with any male in the cluster with the female. t=0 is the moment the roller female is placed on the lawn. Assays were quantified at the three-minute endpoint.

#### Video imaging

Mating pairs (Figure 1B, Videos S1 and S3) were video recorded on a Zeiss AxioObserver.A1 microscope with an Andor iXon3 DU-897 EM-CCD camera operated by MetaMorph 7.7.6 software and a Zeiss Plan-APOCHROMAT 10X/0.32 air objective.

Vulva opening assays (Figure 1C, Video S2, Table 1) were video recorded on a Zeiss Imager.Z1 microscope with an iPhone6/7 camera and LabCam adaptor and a Zeiss EC Plan-NEOFLUAR 20X/0.5 Ph2 air objective; and on a Leica Wild M3Z StereoZoom or a Leica MZ12.5 StereoZoom microscope, with an iPhone6/7 camera and LabCam adaptor (Video S6, Table 1).

Female and male chemotaxis assays (Videos S4 and S5, Figures 2-5, S1-3) were video recorded on a Leica Wild M3Z StereoZoom microscope with an iPhone6/7 camera and LabCam adaptor.

#### Scoring receptive behaviors

Females were scored positive for sedation during a 5-minute assay if they arrested forward and reverse locomotion for at least 30 seconds when the male partner touched his tail to the vulva. Females were scored positive for vulva opening during a 5-minute assay if they visibly contracted the anterior and posterior vulval muscles when the male partner touched his tail to the vulva. Mating success was scored based on visible sperm transfer and/or deposition of a mating plug. For all assays, t=0 is the moment the male begins scanning the female with the ventral side of his tail. For assays of *C. elegans*, *C. briggsae*, and *C. tropicalis* hermaphrodites with conspecific males, each assay included 30 hermaphrodites and one male, and assays were scored manually. For assays of *C. nigoni odr-7* exon1STOP and wild-type controls, each assay included 1-3 females and one male. All other assays in Table 1 included 20 females or hermaphrodites and one male. Sedation and vulva opening were not scored in aged hermaphrodites because they move slowly and have a distended vulva.

#### Reverse Transcription-PCR

Wild-type, exon1STOP, reverted, and exon2STOP *C. nigoni* strains were washed with M9 buffer and extracted for total RNA using TRIzol reagent (Invitrogen) and RNeasy columns (Qiagen). Total cDNAs were generated by reverse transcription with oligo-dT primer (SuperScript III, Invitrogen). *odr-7* cDNAs were amplified by PCR with Q5 polymerase (NEB) and gene-specific primers (see Table S1), visualized on 1% agarose gel, gel-purified (Zymo spin columns) and Sanger sequenced.

#### CRISPR mutagenesis in *C. nigoni*

Inbreeding *C. nigoni* results in sterility, so crosses were designed to retain heterozygosity. We performed *odr-7* CRISPR mutagenesis in *C. nigoni* according to the Mello lab protocol for *C. elegans*^19^ with some modifications. Briefly, virgin 1-day adult wild-type *C. nigoni* females were injected with a CRISPR mix including recombinant Cas9 protein (250 ng/μl), tracrRNA (100 ng/μl), crRNA (56 ng/μl), ssDNA repair oligonucleotide (110 ng/μl), and a *C. elegans rol-6* co-injection marker plasmid (40 ng/μl). See Table S1 for crRNA and ssDNA repair template sequences. Injected females were cultivated at 25-30°C overnight, then bred to wild-type males and plated individually. Plates of F1 larvae and young adults were screened for rollers, and F1 males from roller-containing plates were crossed individually to wild-type females. The F1 males were removed after one day and genotyped for the templated mutation by single-worm lysis (https://bpb-us-w2.wpmucdn.com/sites.wustl.edu/dist/0/1532/files/2018/07/Single-Worm-PCR-r8hwch.pdf), PCR (OneTaq polymerase), 2% agarose gel electrophoresis, gel-purification (Zymo spin columns), and Sanger sequencing. See Table S1 for genotyping PCR primer sequences. F2 heterozygous non-roller females from mutant (X^odr-7^/0) fathers were bred to wild-type males to yield F3 heterozygous and wild-type females and F3 mutant and wild-type males. To reduce inbreeding in the strains, individual F3 male and female animals from different F1 lines were crossed and genotyped. F4 larvae were picked from crosses where the F3 female was heterozygous and the F3 male was mutant. Individual F4 male and female animals from different F3 crosses were crossed and genotyped. F5 mutant lines were identified from crosses where the F4 female was homozygous and the F4 male was mutant. Multiple F5 lines were pooled and verified by genotyping to produce a stable, genetically diverse mutant strain. For the *rol-6* CRISPR mutagenesis, the *rol-6* co-injection marker was omitted and sgRNA (Synthego) was used in place of crRNA and tracrRNA. The dominant *C. elegans* mutation *rol-6(su1006)* provided the sequence used in the template oligonucleotide.^55^ F1 roller males were bred to wild-type females, and four F2 roller males and females were individually crossed to wild-type mates before they were genotyped, confirming the templated mutation (lysis, PCR, purification, and Sanger sequencing). Roller animals were bred from the F3 generation onward to generate a true-breeding roller mutant strain. All *C. nigoni* CRISPR strains were generated from JU1325 except for the *odr-7* reverted strain, which was generated from the *odr-7* exon1STOP strain.

#### DiO staining

1. *C. nigoni* JU1325 or *C. elegans* PD1074 control animals were stained overnight with 40 μg/ml DiO (Molecular Probes) in M9 buffer, then washed with M9 buffer and destained on an OP50 seeded plate for 1 hr before imaging on 2% agarose pads on glass slides. Image stacks (∼20-30 images) were acquired on a Zeiss widefield epifluorescence microscope (Imager.Z1, Axiovision software) with a Zeiss EC Plan-NEOFLUAR 40X/1.3 oil DIC objective. Representative image (Figure S2A) is maximum intensity projection of 17 frames generated using ImageJ software. n = 18 worms imaged.

#### Immunostaining

Wild-type and *odr-7* 3xFLAG *C. nigoni* strains were immunostained according to the Loer lab protocol for serotonin staining in *C. elegans*: http://home.sandiego.edu/~cloer/loerlab/anti5htlong.html. Briefly, worms were washed with M9 buffer, fixed 7 hr in 4% paraformaldehyde on ice, permeabilized 14 hr with 5% beta-mercaptoethanol and 1.5 hr with collagenase, blocked 1 hr in 1% BSA/10% goat serum, incubated overnight at 4°C with primary antibody against FLAG tag and/or serotonin, and incubated 2 hr with green and/or red fluorescently labeled secondary antibody. Antibody dilutions were pre-cleared 1 hr on fixed, permeabilized wild-type *C. nigoni* worms and passed through a 0.22 μm filter prior to use. Antibody concentrations: mouse anti-FLAG, 1:1000; rabbit anti-serotonin, 1:100; goat anti-mouse-Alexa Fluor 488, 1:1000; goat anti-rabbit-TRITC, 1:100.

Stained worms were mounted in VectaShield mounting media with DAPI on 2% agarose pads on glass slides. Image stacks (∼25-30 images) were acquired on a Zeiss widefield epifluorescence microscope (Imager.Z1, Axiovision software) with a Zeiss Plan-APOCHROMAT 63X/1.4 oil objective. Representative images (Figure 4B, Figure S2C-E) are maximum intensity projections of 3-4 frames generated using ImageJ software. n>20 worms imaged for each staining protocol, from three independent batches.

### STATISTICAL ANALYSIS

#### Statistical tests

n, measures of dispersion, and statistical tests for the chemotaxis experiments are reported in the figure captions. n is the number of assays, which each included 20 test animals and one roller target animal. Non-parametric tests were used to compare distributions of females or males in the chemotaxis assays with no assumption of normality. For pairwise comparisons, the Mann-Whitney test was used. For three or more groups, the Kruskal-Wallace test with Dunn’s multiple comparison test was used. Tests were performed with Prism9.2 (GraphPad). In all figures, ns, not significant. * P<0.05, ** P<0.01, *** P<0.001, **** P<0.0001. For within-species comparisons of sedation, vulva opening, and mating success in Table 1, Fisher’s exact test was performed (GraphPad software).

## Supporting information

Supplementary Data Tables for all figures

## Acknowledgments

We thank members of the Bargmann lab, Rene Garcia, Doug Portman, Meital Oren-Suissa, and Patrick Phillips for helpful discussions of this work; Du Cheng for the LabCam adaptor; Max Brown for cultivating strains; Krishna Ghanta and Craig Mello for the CRISPR protocol; Ron Ellis for the *C. briggsae she-1* strain; and Yarden Wiesenfeld for immunostaining protocols and reagents. This work was supported by grants from the Helen Hay Whitney Foundation (to M.S.E.), Howard Hughes Medical Institute (to C.I.B.), and Chan Zuckerberg Initiative (to C.I.B.).

## Author contributions

Investigation, data curation, visualization, writing – original draft, M.S.E. Conceptualization, formal analysis, methodology, resources, validation, writing – review and editing, M.S.E. and C.I.B. Project administration, supervision, funding acquisition, C.I.B.

M.S.E. and C.I.B. conceived the project; M.S.E. performed the experiments, M.S.E. and C.I.B. analyzed the data; M.S.E. and C.I.B. wrote and edited the paper; C.I.B. supervised and funded the work.

## Supplemental Information

Figures S1-S3 and figure legends

Table S1 and table legend

Legends for Videos S1-S6

Supplemental material: A STOP-IN bypass mutant, and references

Supplemental material: *C. nigoni odr-7* cDNA sequences

Key Resources Table

**Figure S1.**
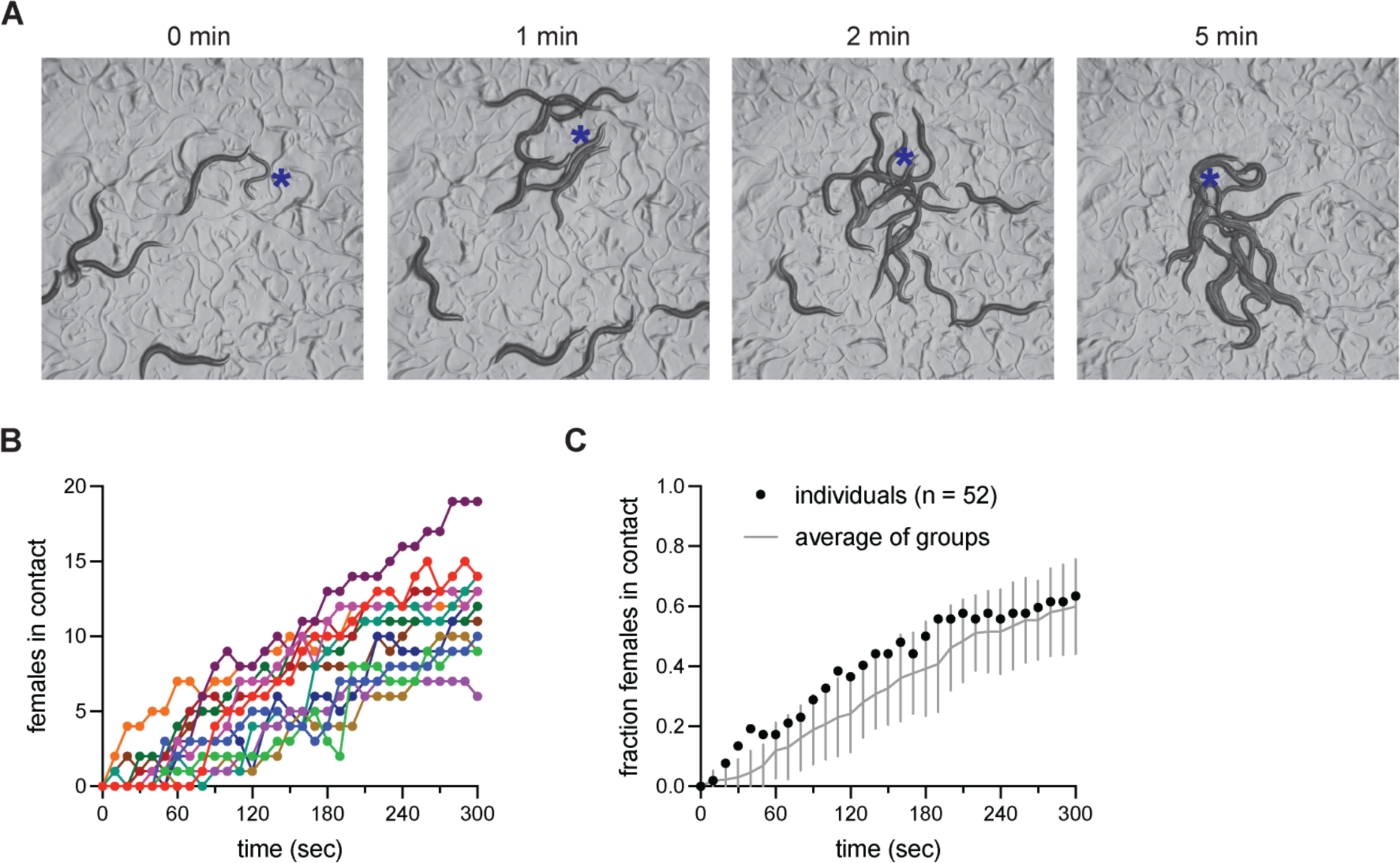
related to Figure 2. Additional chemotaxis assays. (A) Chemotaxis of *C. wallacei* females to *C. wallacei* males. One wild-type target male (blue asterisk) was placed on a lawn with 20 test females. t=0 denotes first male contact with a female. Images are stills from a representative video with times noted. Females are 1 mm long. (B) Results of 13 chemotaxis assays, each with 20 wild-type *C. nigoni* females and one roller *C. nigoni* male, scored over 5 minutes. (C) Results of individual female chemotaxis assays, each with one wild-type *C. nigoni* female and one roller *C. nigoni* male, scored over 5 minutes. (n = 52 assays). Black dots, fraction of females in contact with male at each timepoint. Gray line, average of 13 group assays with 20 females each (Figure S1B); gray bars, +/-SD.

**Figure S2.**
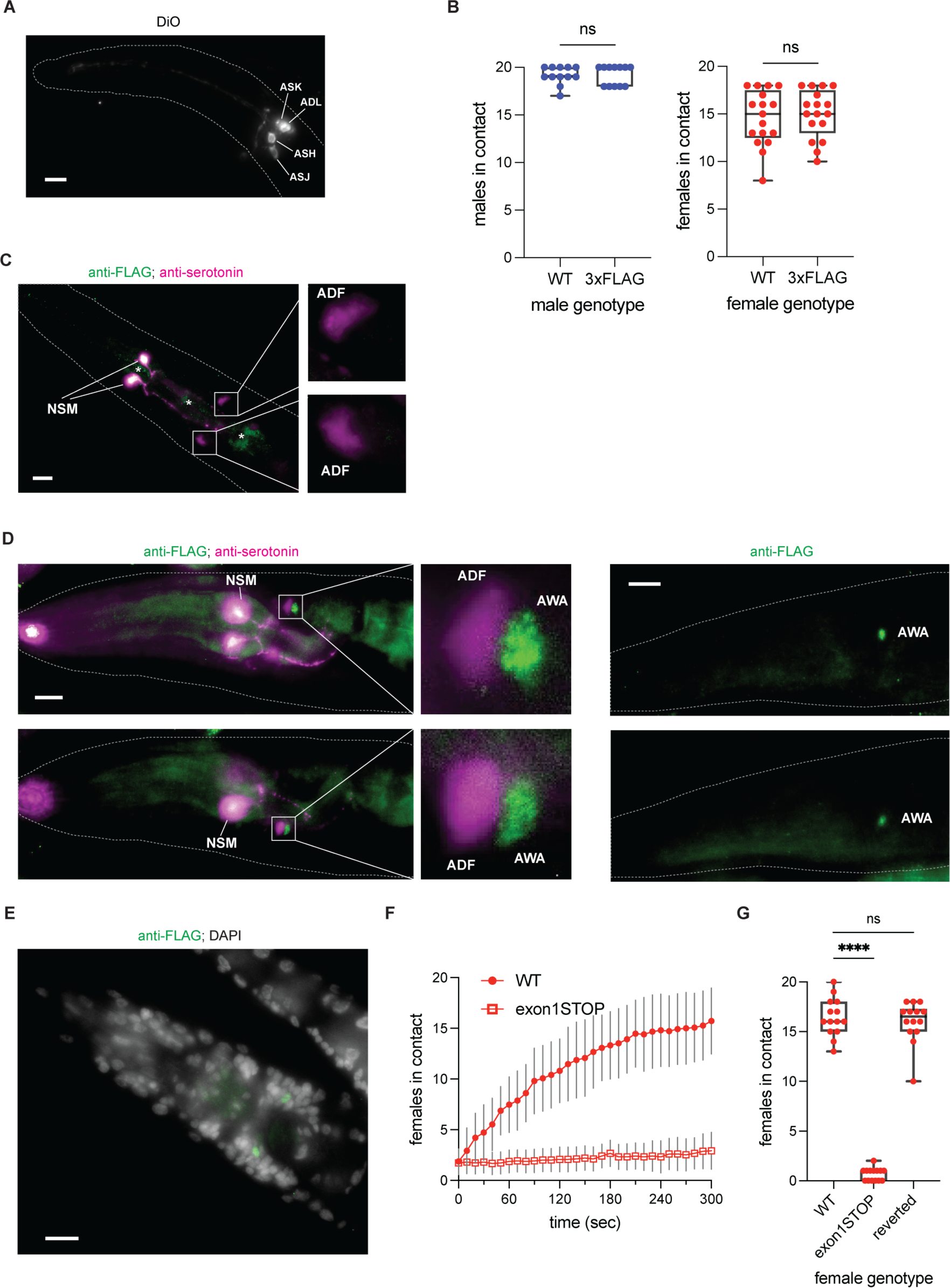
related to Figure 4. Additional analysis of *C. nigoni odr-7*. (A) DiO-stained *C. nigoni*. Neurons are labeled based on similar anatomy to DiO-stained *C. elegans* neurons. Image magnification, 40X. Scale bar, 10 *μ*m. Anterior is at left. (B) Chemotaxis of 20 wild-type or *odr-7::*3xFLAG-tagged *C. nigoni* males to one roller female, scored at three minutes (left, n = 12 assays each) or 20 wild-type or *odr-7::*3xFLAG-tagged *C. nigoni* females to one roller male, scored at five minutes (right, n = 17 assays each). Mann-Whitney test. ns P>0.05. (C) Negative control for immunostaining. Representative wild-type *C. nigoni* adult male imaged for FLAG tag (green) and serotonin (magenta). Note anti-FLAG background staining in the pharynx (asterisks), which is present in other anti-FLAG images as well. (D) ODR-7::FLAG expression in *C. nigoni* females. (Left) Representative adult *odr-7*::3xFLAG female immunostained for FLAG (green) and serotonin (magenta). Inset is magnified in the region of AWA. Labels indicate inferred neuronal identities based on position. (Right) Representative adult *odr-7*::3xFLAG female immunostained for FLAG (green) only. (E) Adult *odr-7*::3xFLAG male immunostained with anti-FLAG antibody (green); staining overlaps with a pair of nuclei in the superimposed DAPI image. In (C-E), image magnification, 63X. Scale bars, 10 *μ*m. Anterior is at left in all images. (F) Chemotaxis of 20 wild-type or *odr-7* exon 1 STOP-IN mutant *C. nigoni* females to one wild-type *C. nigoni* male target (n = 15 assays each). Red dots or squares, mean at each timepoint; gray error bars, +/-SD. (G) Chemotaxis of 20 wild-type, *odr-7* exon 1 STOP-IN mutant, or *odr-7* reverted exon 1 *C. nigoni* females to one roller *C. nigoni* male, with t=0 immediately after female locomotion was stimulated by tail tap; scored at 5 minutes (n = 14 assays each). Kruskal-Wallace test with Dunn’s multiple comparisons test. ns P>0.05, **** P<0.0001.

**Figure S3.**
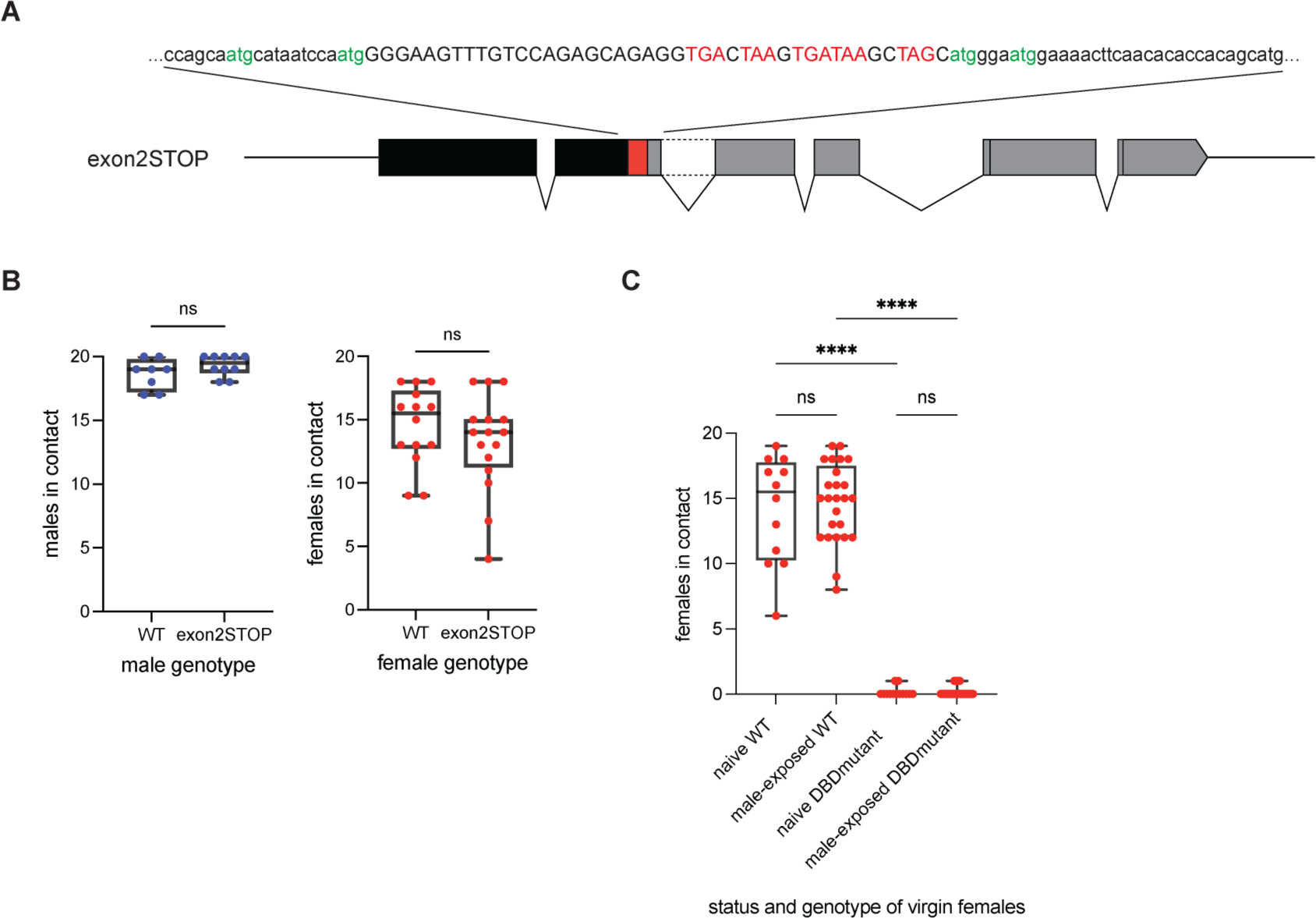
related to Figure 3, Figure 4, Supplemental Information, STAR Methods. An unexpected wild-type phenotype in a *C. nigoni odr-7* mutant. (A) Design of *odr-7* exon 2 STOP-IN CRISPR mutant. Red, 43-bp STOP-IN cassette insertion. Inset sequence labels stop codons (red) and in-frame start codons (green). (B) Chemotaxis of 20 wild-type or *odr-7* exon 2 STOP-IN mutant *C. nigoni* males to a roller female, scored at three minutes (left, n = 8-10 assays each); or *C. nigoni* females to a roller male, scored at five minutes (right, n = 14-16 assays each). Pairwise comparisons: Mann-Whitney test. ns, P>0.05. ****, P<0.0001. (C) Chemotaxis to a roller male of 20 naïve *C. nigoni* females, or 20 virgin *C. nigoni* females that were previously exposed to male odor up to three times (n = 12-29 assays each). Both wild-type and *odr-7* DBD mutants are shown. Naïve and male-exposed data are combined in Figure 4E. Kruskal-Wallace test with Dunn’s multiple comparisons test. ns, P>0.05. ****, P<0.0001.

**Table S1.**
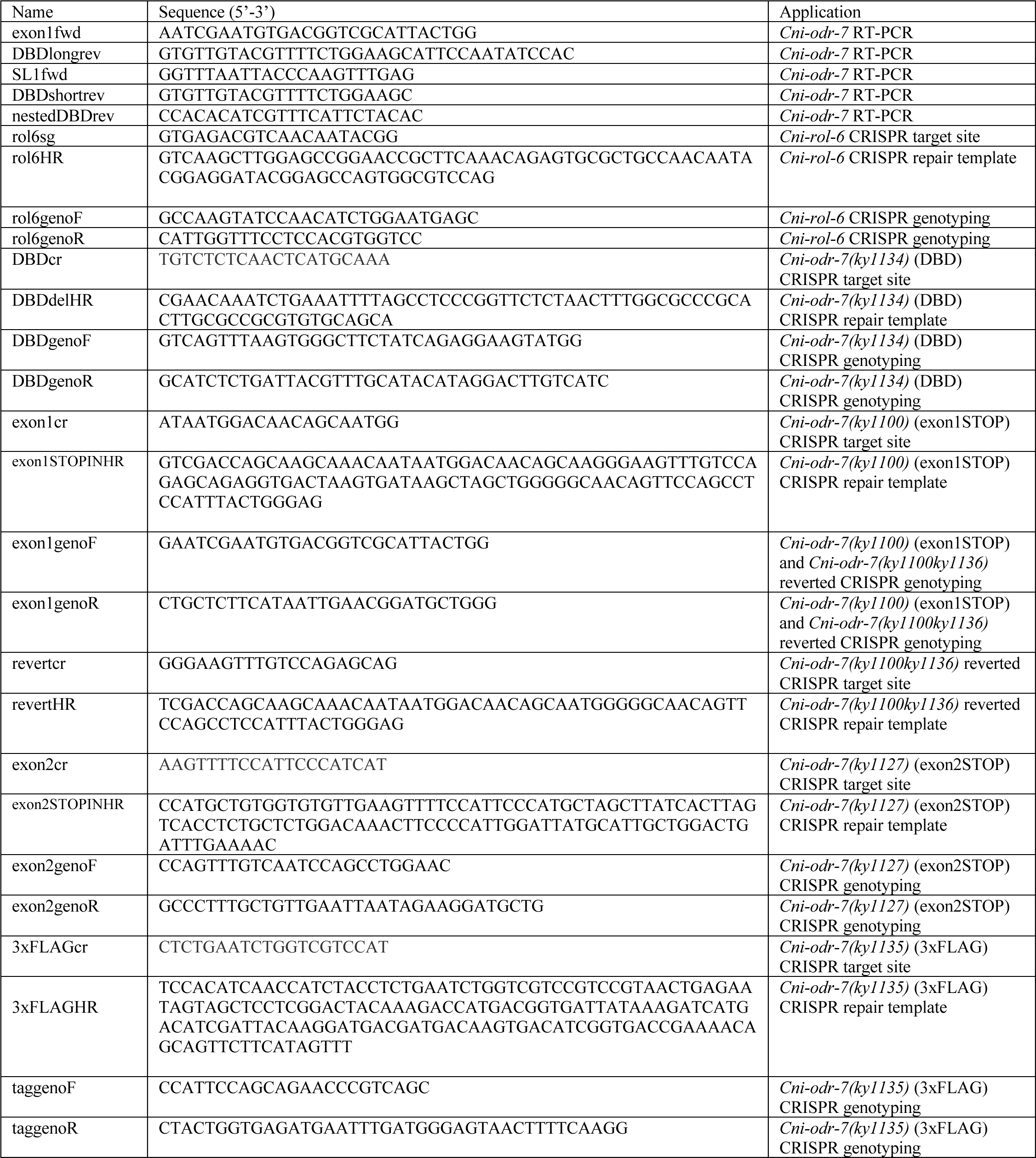
Oligonucleotides.

**Table S1**, related to STAR Methods.

Oligonucleotides. Name, sequence (5’ to 3’), and application are listed.

**Video S1**, related to Figure 1B.

Female sedation behavior in a *C. wallacei* mating pair. The male ventral tail begins scanning the female at 4 s in the video, corresponding to t=0 in Figure 1B. Video speed, 1x.

**Video S2**, Related to Figure 1C.

Female vulva opening behavior in a *C. nigoni* mating pair. The male ventral tail contacts the vulva at 12 s in the video, corresponding to t=0 in Figure 1C. Video speed, 1x.

**Video S3**, related to Figure 1B, 1C. Lack of sedation or vulva opening in a *C. tropicalis* hermaphrodite paired with a *C. wallacei* male. Video speed, 2x.

**Video S4**, related to Figure 2A. *C. nigoni* female chemotaxis to a male. Video of 20 wild-type *C. nigoni* females with one roller *C. nigoni* male added at t=0, recorded for 5 minutes. Video speed, 8x.

**Video S5**, related to Figure 2B. *C. nigoni* male chemotaxis to a female. Video of 20 wild-type *C. nigoni* males with one roller *C. nigoni* female added at t=0, recorded for 3 minutes. Video speed, 4x.

**Video S6**, related to Table 1. Latent sedation and vulva opening in *C. briggsae*. A virgin 1-day adult *she-1 C. briggsae* pseudofemale was partnered with a wild-type *C. nigoni* male.

Pseudofemale is 1 mm long. Video speed, 1x.

## Supplemental Material: A STOP-IN bypass mutant

A *C. nigoni odr-7* mutant (*ky1127*) in which the STOP-IN cassette was inserted into exon 2 (Figure S3A) showed wild-type male and female chemotaxis behavior despite the presence of stop codons in every reading frame (Figure S3B). We considered three possible mechanisms for functional bypass of the STOP-IN cassette: alternative splicing,^S1,S2^ translational readthrough,^S3^ and translational re-initiation.^S4,S5^ Reverse transcription-PCR in the mutant strain did not reveal any exon 2 skipping that could bypass the STOPIN-containing exon at the mRNA level. However, if there were undetected exon 2 skipping, the resulting sequence would be in-frame and could potentially encode functional protein. Translational readthrough is an unlikely mechanism, as it would have to recode two different stop codons in the STOP-IN cassette and correct for a subsequent frameshift that eliminates most of the protein. Translational re-initiation after the STOP-IN cassette could produce a shorter version of ODR-7 that may be functional, as it would contain the DNA binding domain and substantial upstream sequence. The first and third codons immediately downstream of the exon 2 STOP-IN cassette are in-frame start codons (Figure S3A); a similar configuration of stop codons and downstream start codon has been shown to express the downstream open reading frame in *C. elegans*.^S6^ By contrast, the exon 1 mutant’s STOP-IN cassette is not situated near any in-frame start codons; the closest such ATG is 148 bp downstream. The exon 2 STOP-IN result warrants further investigation into the placement of nonsense mutations with respect to in-frame start codons. In line with our results, a similar non-null CRISPR termination codon mutant has been described in *Ooceraea biroi* ants,^S7^ and a study of knockout evasion in a human cell line found residual protein expression in one third of CRISPR frameshift mutations, both from exon skipping and from translational re-initiation.^S8^

## Supplemental Material: *C. nigoni odr-7* cDNA sequences

RT-PCR with an SL1 splice leader forward primer mapped the wild-type *odr-7* mRNA 5’ end to a 29-bp 5’UTR (atttggcctgagatactaacatcaccaga) followed by the WormBase predicted start codon for *Cni-odr-7.1*. Coding transcripts below. Coding exons are separated by commas.

### *C. briggsae*-like splice form

Aligns with WormBase *Cbr-odr-7* coding transcript CBG00123.1. ATGAATAAAGGAATGCTCGGGTATTACTGTATCAATTGTATCATGGCAATGCCACATTTCACTTGTTATCCAATGATTTTATCAGAATCGAATGTGACGGTCGCATTACTGGAAAACATGGCAGAAAAGGTCGACCAGCAAGCAAACAATAATGGACAACAGCAATGGGGGCAACAGTTCCAGCCTCCATTTACTGGGAGACCTCCCGGATCCCAGCATCCGTTCAATTATGAAGAGCAGCACAGAGTCAGAGAACCAATTTCACATGATTTCGATAATGATTTATCCCGACAGCAGACAACAGCCCCATTCATGCAACACTTTTTTCCCAGAATAG,TTC CGAGTTTACCATTTCCCGATTTCACCGACTACCAAAGGTTTAACGGTTTTCAACGTAATGCATTCTTTCCAACTCCTTTCGGTCCCCAGTTTGTCAATCCAGCCTGGAACAGTTTTCAAATCAGTCCAGCAATGCATAATCCAATGATGGGAATGGAAAACTTCAACACACCACAGCATG,CGGAATACAATATCAAGCAGTCCCTGGACCAAAAGTTTGATAGCGGATCGGATATTAAATCGGTTAAAGACGAGCCAATGGAAAACTTGGATAACAAGAAGAAGAATGAAGCCAAGGCAGATCATCAGAGATACTCATCTCTTCCCACAGATAAACAATTTGATCAGTCGAG,AACTGCGCTACCATCATTCACCCCAACATTTTATCCTGTTCCTGTAGCTCCATTAGACATGGCTACGAACACCACTTTCAAGCAGGAATTGAGCCCG,CCTCCCGGTTCTCTACACGATTGCCAGGTCTGTCTCTCAACTCATGCAAATGGTCTTCACTTTGGCGCCCGCACTTGCGCCGCGTGTGCAGCATTTTTCAGAAGGACAATCAGCGATGACAAGTCCTATGTATGCAAACGTAATCAGAGATGCACAAACGCTAGTCGTGATGGTACTGGATATCGGAAGATTTGCCGTGCGTGTAGAATGAAACGATGTGTGGATATTGGAATGCTTCCAGAAA,ACGTACAACACAAGCGGGCGCGAAGAGAATCCACCGGCTCCACACCTCCTCCGCCGAAAATTGGGTTCGATAACTTCTTCCCTGGTTGTTTCTATCCGCCATTCCAGCAGAACCCGTCAACCGTTCCACATCAACCATCTACCTCTGAATCTGGTCGTCCATCGGTGACTGAAAACAACAGTTCTTCATAG

### Intron-retaining splice form

WormBase *Cni-odr-7.1* coding transcript. The retained intron is underlined. ATGAATAAAGGAATGCTCGGGTATTACTGTATCAATTGTATCATGGCAATGCCACATTTCACTTGTTATCCAATGATTTTATCAGAATCGAATGTGACGGTCGCATTACTGGAAAACATGGCAGAAAAGGTCGACCAGCAAGCAAACAATAATGGACAACAGCAATGGGGGCAACAGTTCCAGCCTCCATTTACTGGGAGACCTCCCGGATCCCAGCATCCGTTCAATTATGAAGAGCAGCACAGAGTCAGAGAACCAATTTCACATGATTTCGATAATGATTTATCCCGACAGCAGACAACAGCCCCATTCATGCAACACTTTTTTCCCAGAATAG,TTC CGAGTTTACCATTTCCCGATTTCACCGACTACCAAAGGTTTAACGGTTTTCAACGTAATGCATTCTTTC CAACTCCTTTCGGTCCCCAGTTTGTCAATCCAGCCTGGAACAGTTTTCAAATCAGTCCAGCAATGCATA ATCCAATGATGGGAATGGAAAACTTCAACACACCACAGCATG<u>GTCAGCATCCTTCTATTAATTCAACA GCAAAGGGCAAAGAAAGTCTTCAGTCTTATCAAGGGACACCTTTGACTCAAGCTACTGATAGCACCCC AGATAAACCACCAGTTCTTTTAG</u>CGGAATACAATATCAAGCAGTCCCTGGACCAAAAGTTTGATAGCG GATCGGATATTAAATCGGTTAAAGACGAGCCAATGGAAAACTTGGATAACAAGAAGAAGAATGAAGCCAAGGCAGATCATCAGAGATACTCATCTCTTCCCACAGATAAACAATTTGATCAGTCGAG,AACTGCGC TACCATCATTCACCCCAACATTTTATCCTGTTCCTGTAGCTCCATTAGACATGGCTACGAACACCACTTTCAAGCAGGAATTGAGCCCG,CCTCCCGGTTCTCTACACGATTGCCAGGTCTGTCTCTCAACTCATGCAAATGGTCTTCACTTTGGCGCCCGCACTTGCGCCGCGTGTGCAGCATTTTTCAGAAGGACAATCAGCGATGACAAGTCCTATGTATGCAAACGTAATCAGAGATGCACAAACGCTAGTCGTGATGGTACTGGATATCGGAAGATTTGCCGTGCGTGTAGAATGAAACGATGTGTGGATATTGGAATGCTTCCAGAAA,ACGTACAA CACAAGCGGGCGCGAAGAGAATCCACCGGCTCCACACCTCCTCCGCCGAAAATTGGGTTCGATAACTTCTTCCCTGGTTGTTTCTATCCGCCATTCCAGCAGAACCCGTCAACCGTTCCACATCAACCATCTACCTCTGAATCTGGTCGTCCATCGGTGACTGAAAACAACAGTTCTTCATAG

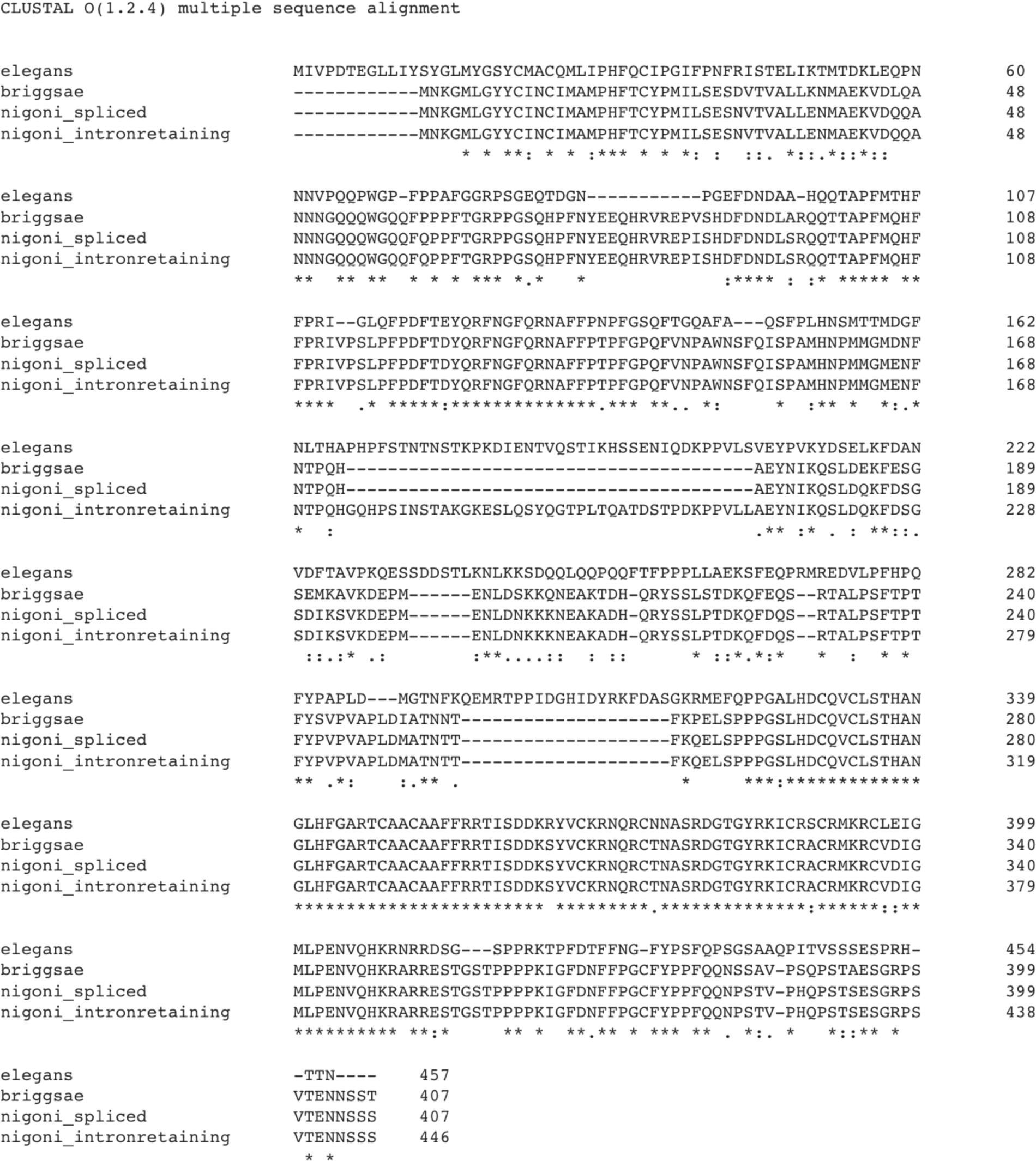

## Key resources table

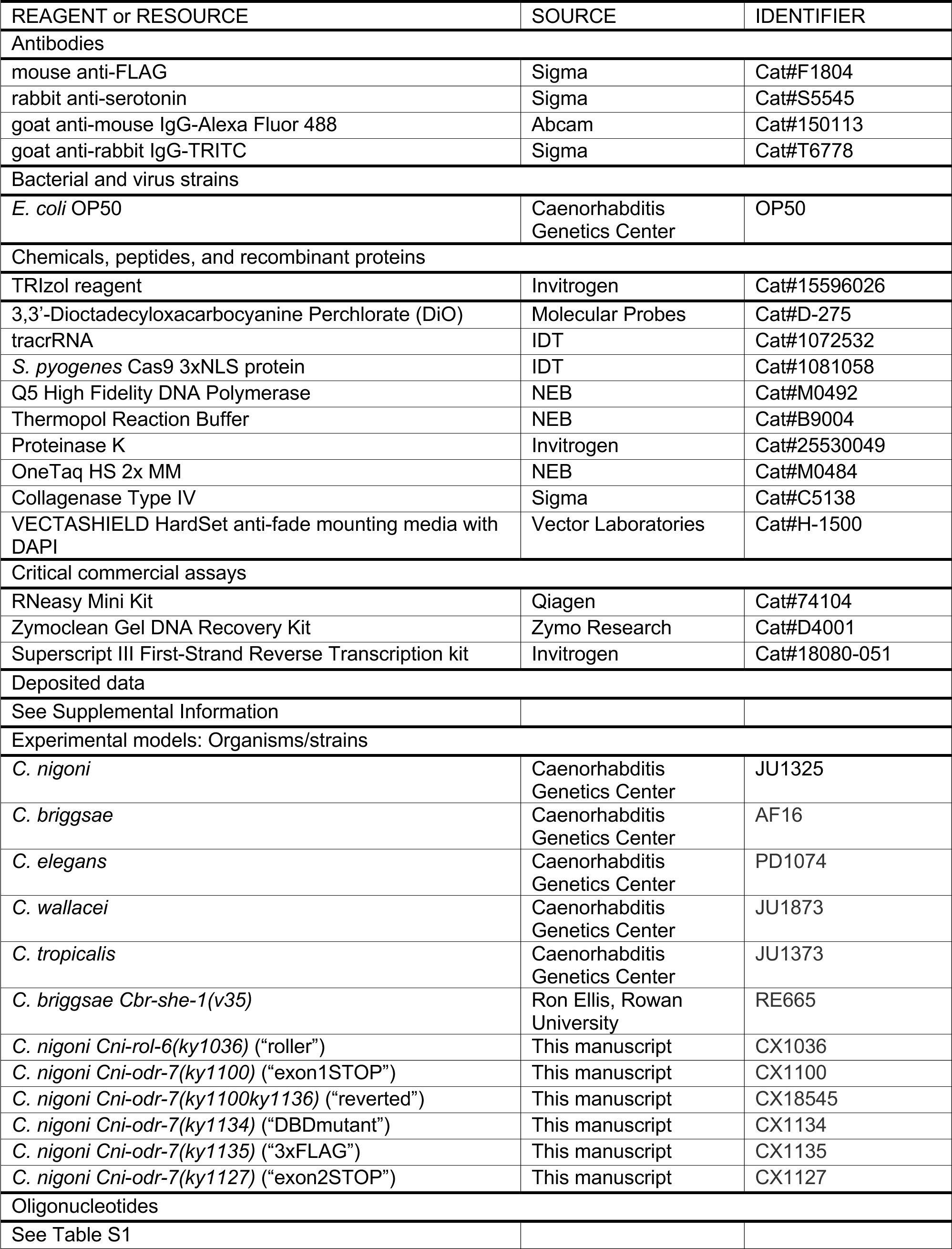
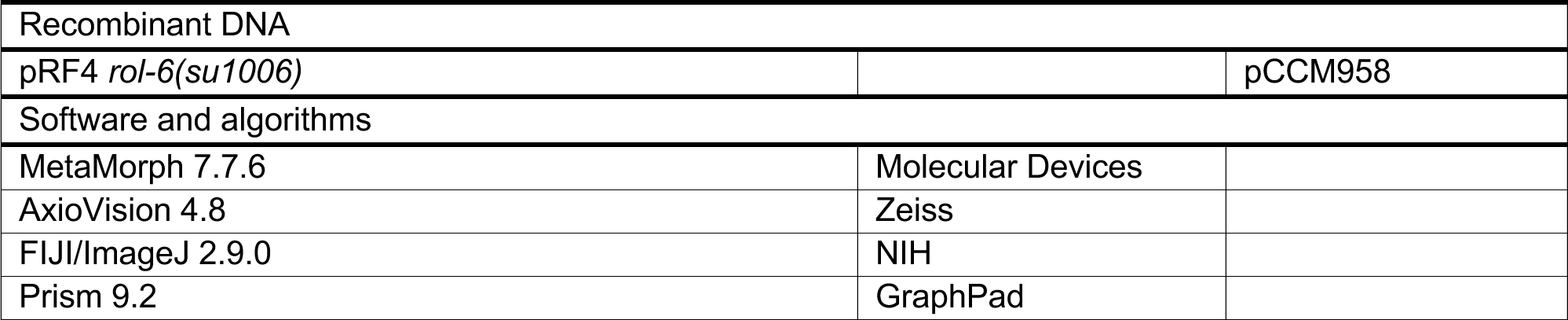

